# Primary auditory thalamus relays directly to cortical layer 1 interneurons

**DOI:** 10.1101/2024.07.16.603741

**Authors:** Lucas G. Vattino, Cathryn P. MacGregor, Christine Junhui Liu, Carolyn G. Sweeney, Anne E. Takesian

## Abstract

Inhibitory interneurons within cortical layer 1 (L1-INs) integrate inputs from diverse brain regions to modulate sensory processing and plasticity, but the sensory inputs that recruit these interneurons have not been identified. Here we used monosynaptic retrograde tracing and whole-cell electrophysiology to characterize the thalamic inputs onto two major subpopulations of L1-INs in the mouse auditory cortex. We find that the vast majority of auditory thalamic inputs to these L1-INs unexpectedly arise from the ventral subdivision of the medial geniculate body (MGBv), the tonotopically-organized primary auditory thalamus. Moreover, these interneurons receive robust functional monosynaptic MGBv inputs that are comparable to those recorded in the L4 excitatory pyramidal neurons. Our findings identify a direct pathway from the primary auditory thalamus to the L1-INs, suggesting that these interneurons are uniquely positioned to integrate thalamic inputs conveying precise sensory information with top-down inputs carrying information about brain states and learned associations.

## Introduction

Our perception of the external world is powerfully influenced by internal states and learned associations. Given that our acoustic environment is generally a chaos of sounds, accurate perception requires that the auditory circuits dynamically allocate resources to process only meaningful information. For this, the auditory system relies on the convergence of sound information with signals that convey contextual information such as vigilance, locomotion, and the acquisition of rewards^1–3^. In the auditory cortex (ACx), sound processing relies on the coordinated activity of excitatory and inhibitory neurons to represent the spectro-temporal structure of sounds from signals arriving from the auditory thalamus, the medial geniculate body (MGB)^4–6^. These ascending inputs are integrated with an elaborate network of top-down and neuromodulatory projections that filter and modify the auditory signals, acutely influencing sound perception and sound-driven behaviors^7–18^. Moreover, the convergence of these top-down and neuromodulatory inputs with sensory signals can induce long-lasting changes in ACx that may underlie auditory learning^7,19–30^. However, where these diverse inputs converge within the complex ACx circuits is not well understood.

Across sensory cortices, the outermost cortical layer (layer 1, L1) is densely populated by axons arising from neuromodulatory centers^9,21,24,31–35^, other sensory, motor, and associative cortices^36–44^, as well as limbic regions^44–46^. L1 in sensory cortices is also characterized by a dense lattice of axons arising from higher-order thalamic nuclei^47–54^. Within the primary auditory cortex (A1), axons from the higher-order auditory thalamus, the medial (MGBm) and dorsal (MGBd) divisions of the MGB, predominantly target L1^51,52^. These higher-order thalamic neurons receive inputs from auditory midbrain, thalamic and cortical areas, as well as from neuromodulatory and integrative centers throughout the brain, and are thought to convey multimodal sensory and non-sensory information to the ACx^47,55,56^. There are also a number of intriguing reports across sensory systems that axons originating in the primary thalamic nuclei, thought to primarily target cortical L4, also extend to L1^24,44,46,49,51,57–59^. These nuclei relay fast sensory signals that are tuned to specific features of auditory, visual, and somatosensory stimuli. For example, the primary auditory thalamus, the ventral subdivision of the MGB (MGBv), receives ascending inputs from the central nucleus of the inferior colliculus and relays fast, frequency-tuned sound information to A1^5,56,60^. MGBv axons are known to densely innervate L3b and L4 but have been recently shown to extend to L1^24,51,57,59^. Moreover, the thin band of MGBv axons within L1 in ACx is frequency-tuned and may form a coarse topographic map of sound frequency representation that mirrors the tonotopic map in L4^24,51^. Therefore, across sensory modalities, L1 is a site where diverse multimodal, neuromodulatory, top-down and sensory inputs may be integrated^34,61,62^, but the cellular targets of these inputs have not been fully identified.

L1 is populated by sparse GABAergic inhibitory INs that can be subdivided into two major subtypes based on the expression of the molecular markers vasoactive intestinal peptide (VIP) and neuron-derived neurotrophic factor (NDNF)^63^. Interest in these L1-INs has recently intensified as several studies have revealed that they powerfully modulate cortical activity and underlie context-dependent behavior and learning^8,21,39,44,47,64–67^. VIP- and NDNF-INs robustly express neuromodulatory receptors^68–71^ and receive inputs from neuromodulatory regions, such as the cholinergic basal forebrain^39,44,72^. In the ACx, L1-INs also receive inputs from other sensory cortices, as well as top-down inputs from cortical association areas such as the prefrontal, retrosplenial and insular cortices, among others^39^. Therefore, L1-INs have been recently highlighted as mediators of top-down inputs that broadcast information about global brain states and contextual cues across cortex^34,73^. However, an alternative model of these L1-INs is that they are activated by specific sensory inputs inherited by the axons from the primary thalamic nuclei that arborize in L1^24,34,51^. This circuit organization would place L1-INs in a unique position to integrate neuromodulatory and top-down inputs with inputs relaying feature-specific sensory information^34,62^. This model is consistent with the recent evidence that L1-INs send spatially precise projections to their cortical targets^24,39,74,75^, punching ‘holes’ in local inhibitory networks to modulate the gain of cortical columns^44,67^. For example, small subsets of L1-INs in ACx within cortical frequency columns may be recruited by specific sounds signaled by the axons from the primary MGBv. In turn, these L1-INs may activate cortical columns to tune the cortical neurons below^21,24,39,44,75^, via both narrowly descending projections that target other INs and horizontal projections to pyramidal neuron dendrites^24,63,75^. Thus, identifying the sources of thalamic input to L1-IN subtypes will provide important insight into their function within the cortical network.

Here, we mapped the connectivity between the auditory thalamus and the two major L1-IN subtypes and characterized the electrophysiological properties of the thalamocortical synapses onto these INs. Unexpectedly, we found that L1-INs subtypes receive the vast majority of their auditory thalamic inputs from the primary MGBv. Moreover, we found that the MGBv neurons form strong monosynaptic connections with L1 VIP- and NDNF-INs in A1, comparable in strength, kinetics, and short-term plasticity to the thalamocortical synapses onto L4 excitatory pyramidal (L4 Pyr) neurons. Together, our results identify a direct relay pathway between the MGBv and L1 VIP- and NDNF-INs in ACx, suggesting that these interneurons process information about the spectro-temporal features of sounds inherited from this primary auditory thalamus. These findings point to L1-INs as sites within the complex cortical circuitry where precise auditory information and neuromodulatory signals converge to modulate deeper-layer ACx neurons in a sound-specific manner. Together, these results contribute to our understanding of the specific inputs that recruit L1-INs, providing new insight into the auditory cortical circuits that mediate context-dependent sensory processing and learning.

## Results

### VIP- and NDNF-INs in superficial ACx receive the majority of thalamic inputs from the MGBv

The auditory thalamus is composed of several distinct subnuclei that receive inputs from diverse brain regions^56^ and relay both sensory and non-sensory information to the ACx. To investigate the connectivity between these thalamic subnuclei and VIP- and NDNF-INs in superficial ACx, we used rabies virus-based monosynaptic retrograde tracing^76^ (**Figure 1A**). We restricted rabies virus infection to VIP- and NDNF-INs by injecting a Cre-dependent adeno-associated viral vector (AAV) encoding the avian EnvA receptor, TVA, fused to mCherry into superficial ACx of adult Vip-IRES-Cre or Ndnf-IRES-Cre mice. We also co-injected a Cre-dependent AAV encoding the rabies virus glycoprotein (RVG) to allow for retrograde transfer of RVG-deleted rabies virus. To restrict our starter cell population to the superficial cortical layers, we injected these viruses just below the pial surface. Two weeks later, the mice were injected in the same site with EnvA-pseudotyped RVG-deleted rabies virus encoding nuclear-localized EGFP (RV-nEGFP). Coronal brain sections were collected one week following the RV-nEGFP injection, and the 3-D brain volume containing all the infected starter cells and the auditory thalamus was imaged and reconstructed. We used semi-automatic cell counting to quantify the number and distribution of infected starter cells within ACx and putative presynaptic neurons within auditory thalamic regions.

**Figure 1.**
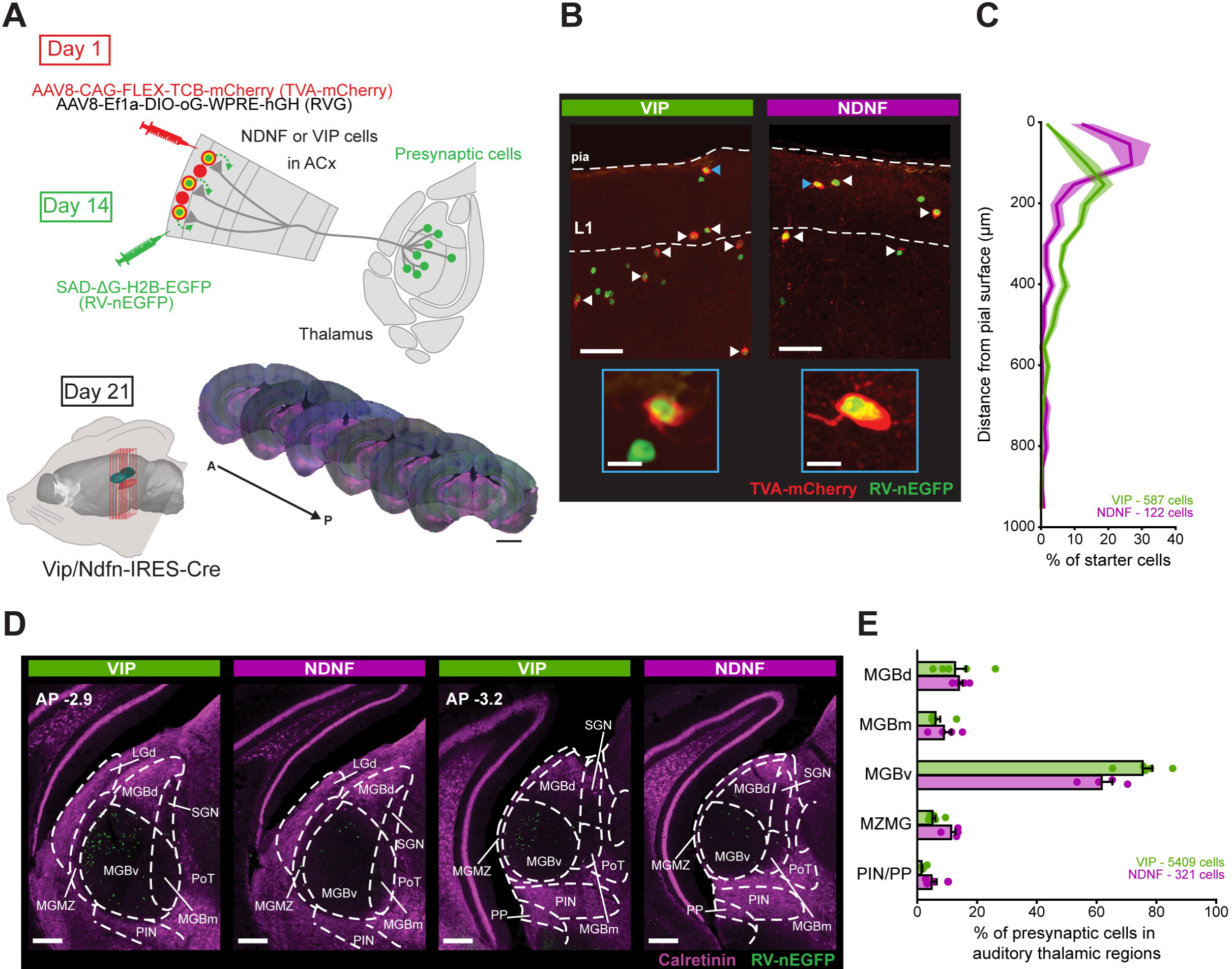
VIP- and NDNF-INs in ACx receive monosynaptic inputs from diverse auditory thalamic subnuclei. **(A)** Schematic of the experimental procedure to identify presynaptic neurons in thalamus that synapse onto VIP- and NDNF-INs. Mice were injected in the left ACx with Cre-dependent viruses expressing the avian EnvA receptor (TVA-mCherry) and the rabies virus glycoprotein (RVG). Two weeks later, a constitutive RVG-deleted, EnvA-pseudotyped rabies virus expressing nuclear-targeted EGFP (RV-nEGFP) was injected into the same sites. The TVA allows for RV-nEGFP to infect starter cells, and the RVG enables retrograde monosynaptic infection of presynaptic partners. Following one week, brain sections (50 µm) were collected, incubated with DAPI and anti-calretinin to delineate the higher-order auditory thalamic regions. Sections were then imaged and aligned to a common coordinate framework atlas for posterior starter and presynaptic cell counting. Scale bar: 2 mm. **(B)** Example confocal images of starter cells (arrows) in ACx of Vip-IRES-Cre and Ndnf-IRES-Cre mice. Starter cells are identified by the co-expression of TVA-mCherry and RV-nEGFP. Scale bar: 50 µm. Inset: Detail of the starter cells indicated with blue arrows. Scale bar: 10 µm. **(C)** Distribution of starter cells across cortical depth. Numbers of total starter cells in auditory cortical areas (A1, AuD, AuV and TeA) are indicated for each genotype. **(D)** Example epifluorescent images of presynaptic RV-nEGFP^+^ cells at two different anterior-posterior (AP) coordinates encompassing the auditory thalamic regions. Calretinin immunostaining delineates the higher-order auditory thalamic areas, whereas the region showing no calretinin staining corresponds to the primary MGBv. **(E)** Distribution of the density of RV-nEGFP^+^ presynaptic cells within auditory thalamic regions. Each data point represents an individual mouse. A1: primary ACx; AuD: auditory dorsal area; AuV: auditory ventral area; TeA: temporal association cortex; MGBv: ventral division of the medial geniculate body; MGBm: medial division of the medial geniculate body; MGBd: dorsal division of the medial geniculate body; MZMG: marginal zone of the medial geniculate body; PIN: posterior intralaminar nucleus; PP: peripeduncular nucleus.

Starter cells were identified by the co-expression of RV-nEGFP and TVA-mCherry (**Figure 1B** and **Supplementary Figure 1** - controls for the specificity of TVA-mCherry to Cre-expressing neurons are included in **Supplementary Figure 2**A-C). The majority of NDNF-IN starter cells were located close to the pial surface whereas the majority of VIP-IN starter cells were located along the L1/L2 border (**Figure 1B**,**C** and **Supplementary Figure 1**), consistent with the distribution of these cortical INs shown in previous studies^63,77^. The majority of infected starter cells were located in A1, but we found some starter cells in abutting regions (**Supplementary Figure 1**). Putative presynaptic neurons were identified by the expression of only RV-nEGFP (**Figure 1D** - controls showing the requirement of RVG for monosynaptic transfer of RV-nEGFP are included in **Supplementary Figure 2**D-H). To delineate the MGB subdivisions, we aligned the brain sections to a unified common coordinate framework atlas (YSK atlas)^78^ in combination with immunostaining against calretinin, a marker for higher-order auditory thalamic regions^51,52,79^ (**Figure 1D**).

Consistent with previous anatomical studies mapping thalamic inputs to the ACx, we found a robust source of presynaptic neurons in the ipsilateral MGB^80^. We found a scattering of presynaptic neurons to VIP- and NDNF-INs located in the higher-order auditory thalamus, in line with anatomical studies showing strong innervation of superficial ACx by these nuclei^47,51,52,59^ (**Figure 1D**,**E**). Strikingly, however, the largest fraction of presynaptic neurons within auditory thalamus to both superficial IN subtypes was located within the primary subdivision of the MGB, the MGBv (**Figure 1D**,**E**). We found just modest differences between VIP- and NDNF-INs in the distribution of presynaptic neurons across the distinct auditory thalamic subregions, with a slightly higher fraction of higher-order thalamic inputs to NDNF-INs (**Supplementary Table 1**). However, these results could be due to small differences in the distribution of starter cells across auditory cortical areas (**Supplementary Figure 1**). Together, our findings suggest that both VIP- and NDNF-INs within superficial ACx receive monosynaptic connections from neurons arising from several distinct subdivisions of the auditory thalamus; however, the vast majority of presynaptic neurons are concentrated in the primary MGBv.

### MGBv neurons form functional monosynaptic connections with L1 VIP- and NDNF-INs in ACx

To confirm that the MGBv neurons form direct, functional synapses onto the L1-INs in ACx, we performed whole-cell voltage clamp recordings from L1 VIP-INs, L1 NDNF-INs and L4 pyramidal excitatory (L4 Pyr) neurons in the same slices while optogenetically activating MGBv afferents. We labeled these L1-INs by injecting AAVs encoding Cre-dependent TdTomato and Flp-dependent mNeonGreen within the ACx of adult Vip-IRES-Cre x Ndnf-IRES-FlpO mice. In the same procedure, we injected an AAV encoding ChR2 fused to mCherry within the MGBv of these mice (**Figure 2A**, **left**). We observed strong bands of mCherry-labeled MGBv axon terminations within L4 and L1 of ACx (**Figure 2A**, **right**), as shown previously^24,51,81^. To characterize the thalamocortical synapses onto the L1-IN subtypes, we obtained optogenetically-evoked excitatory postsynaptic currents (EPSCs) from L1 VIP-INs, L1 NDNF-INs, and L4 L4 Pyr neurons in A1 by recording at a holding potential (V_hold_) of -70 mV. Cells were filled with biocytin for post-hoc anatomical reconstructions (**Figure 2B**,**C**). We found robust light-evoked EPSCs in all three cell types that we confirmed to be monosynaptic by blockade with the voltage-gated Na^+^ channel blocker TTX (1 µM) and rescue with the voltage-gated K^+^ channel blocker 4-AP (1 mM)^82,83^ (**Figure 2D**). In the presence of TTX and 4-AP, all three cell types exhibited short EPSC onset latencies, consistent with optogenetically-activated monosynaptic responses (**Figure 2E** and **Supplementary Table 2**). We further confirmed that these monosynaptic responses were mediated by glutamate receptors by the elimination of EPSCs with the AMPA receptor blocker DNQX (20 µM) and the NMDA receptor blocker AP5 (50 µM) (**Figure 2D**). Surprisingly, we found no differences in monosynaptic EPSC amplitudes among L1 VIP-INs, L1 NDNF-INs, and L4 Pyr neurons (**Figure 2E** and **Supplementary Table 2**). The response magnitude depends on the number of stimulated axons, which may vary across slices with differences in ChR2 viral expression. To control for these potential differences, monosynaptic EPSCs in L1-INs were normalized to those recorded in L4 Pyr within the same slice. We found no significant differences in the normalized amplitudes across the three cell types (**Figure 2E** and **Supplementary Table 2**). Together, our results indicate that both L1 VIP- and NDNF-INs in ACx receive functional monosynaptic inputs from the MGBv, and that the amplitudes of these EPSCs in L1-INs are comparable to those recorded in L4 Pyr neurons.

**Figure 2.**
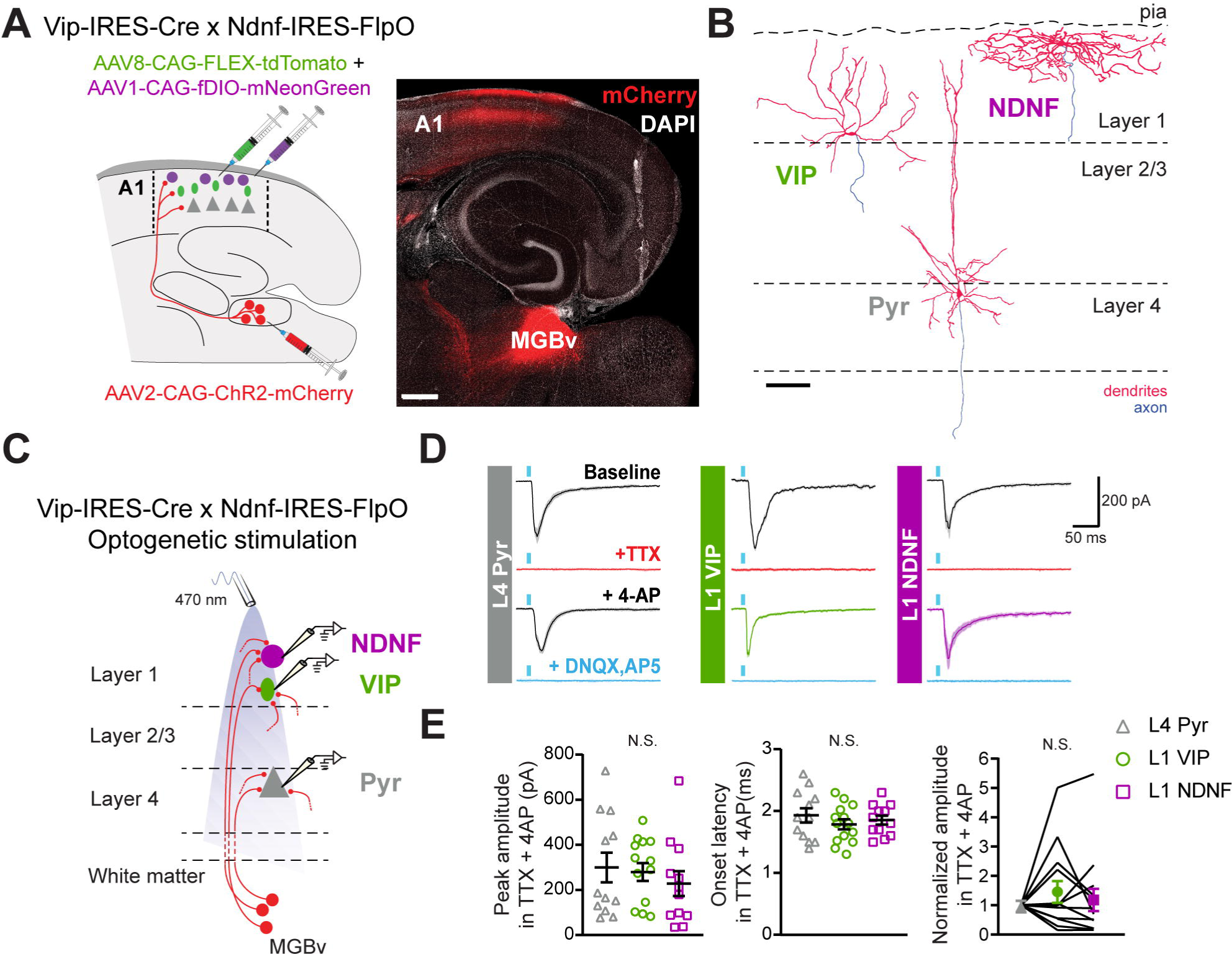
VIP- and NDNF-INs in layer 1 of ACx receive functional monosynaptic inputs from the ventral portion of the MGB (MGBv). **(A) Left.** Schematic of the experimental procedure to obtain ChR2-mediated monosynaptic currents in layer 1 (L1) VIP-INs (green), L1 NDNF-INs (magenta) and layer 4 pyramidal excitatory (L4 Pyr; gray) neurons within thalamocortical slices. **Right.** Confocal image of a representative thalamocortical slice showing mCherry-ChR2 expression in the ventral division of the medial geniculate body (MGBv) and thalamocortical axons within layers L1 and L4 of primary auditory cortex (A1). Scale bar: 500 µm. **(B)** Reconstruction of recorded L4 Pyr, L1 VIP- and L1 NDNF-INs (). Scale bar: 100 µm. **(C)** Schematic of the recording conditions to obtain light-evoked EPSCs from L4 Pyr, L1 VIP- and L1 NDNF-INs by activating ChR2-expressing MGBv axons. Recordings from at least one neuron of each type were obtained in every slice. **(D)** Example voltage clamp recordings (V_hold_ = -70 mV; mean ± SD of 10 trials) showing the effects of sequential addition of TTX and 4-AP on light-evoked EPSCs. Currents were abolished by glutamatergic receptor blockers DNQX and AP5. **(E)** Peak amplitude (**left**) and onset latency (**middle**) of monosynaptic EPSCs recorded in TTX and 4-AP from the three cell types. EPSC amplitudes normalized to the average EPSC amplitude of the L4 Pyr neurons recorded in the same slice (**right**). See **Supplementary Table 2** for details on statistical tests. N.S., p > 0.05.

### L1-INs and L4 Pyr neurons show comparable thalamic-evoked monosynaptic responses

While we relied on optogenetic experiments to detect monosynaptic connections between the MGBv and L1-INs, the use of AAV vectors, ChR2, TTX, and 4-AP can influence the synaptic response properties^84,85^. We therefore used electrical stimulation of thalamic axons to compare the properties of thalamocortical synapses onto L1 VIP-INs, L1 NDNF-INs and L4 Pyr neurons in A1. We cut thalamocortical slices from Vip-IRES-Cre x Ai9 or Ndnf-IRES-Cre x Ai9 mice, which express TdTomato in either VIP- or NDNF-INs. The peri-horizontal thalamocortical slice^24,60,86^ preserves the axons between the MGBv and cortex, allowing for the direct electrical stimulation of thalamocortical axons using a bipolar electrode (). We first studied the EPSCs (V_hold_ = -70 mV) evoked by electrical activation of putative single thalamic axons using minimal stimulation (**Figure 3A**;)^87,88^. We found that minimal stimulation-evoked EPSC amplitudes in L1 VIP- and NDNF-INs were comparable to those recorded in L4 Pyr neurons (**Figure 3B**,**C** and **Supplementary Table 2**). The onset latencies of putative single thalamic axon responses were significantly longer for the L1-INs as compared to the L4 Pyr neurons, reflecting an expected transmission delay of thalamic axons between L4 and L1^24,89^ (**Figure 3C** and **Supplementary Table 2**). However, the three cell types showed comparable rise times, measured as the time between the onset of the response and the peak of the EPSC (**Figure 3C** and **Supplementary Table 2**). These data show that L1 VIP- and NDNF-INs in A1 show responses to activation of putative single thalamic axons that are comparable to those observed in L4 Pyr neurons.

**Figure 3.**
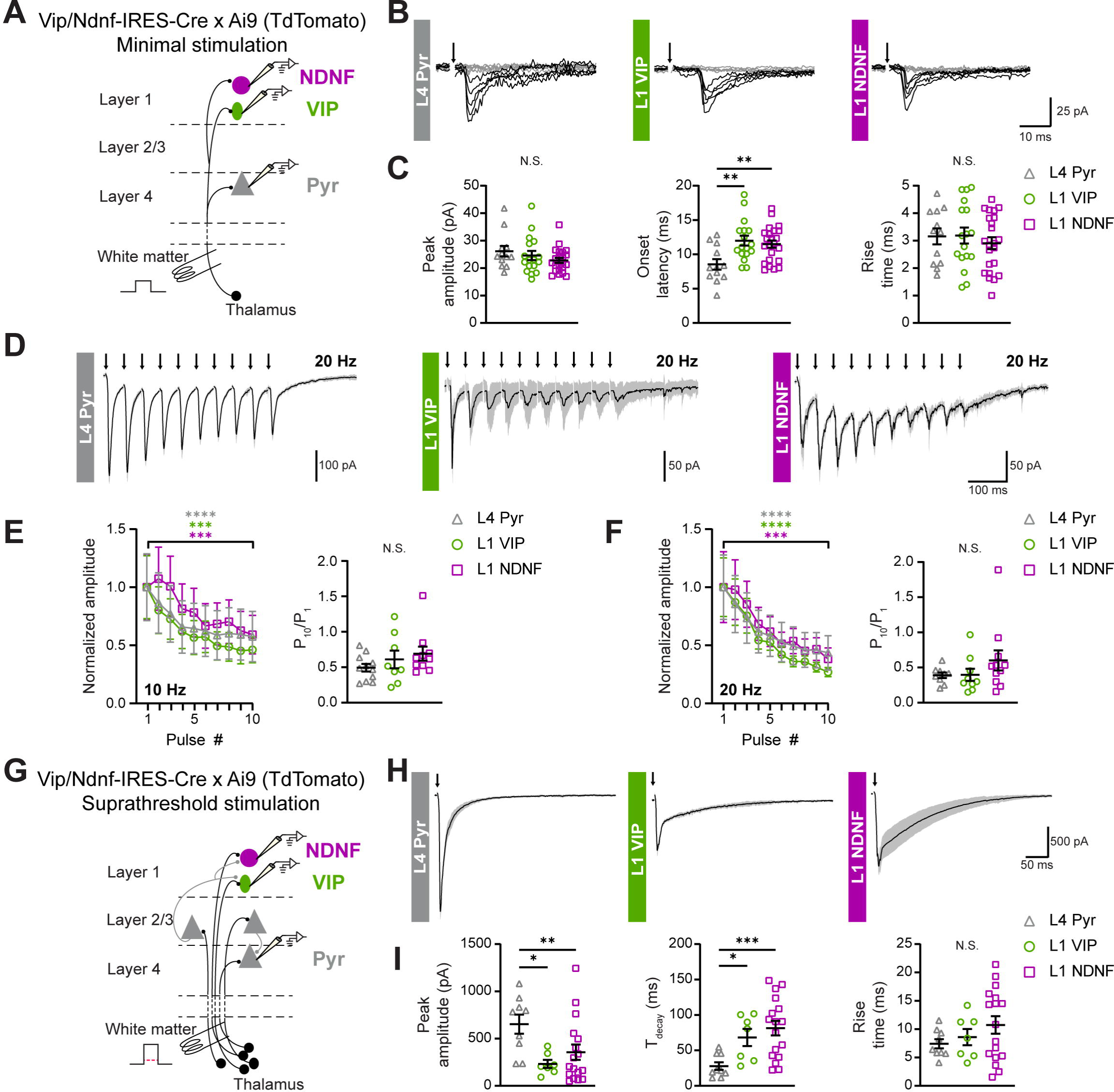
Synaptic properties of thalamocortical inputs onto L4 Pyr, L1 VIP-INs and L1 NDNF-INs. **(A)** Schematic of the experimental procedure to record EPSCs electrically-evoked by minimal stimulation of the thalamic axons within thalamocortical slices. The stimulus intensity was adjusted to obtain a 50% failure rate. Minimal stimulation-evoked EPSCs were recorded from L1 VIP- and NDNF-INs in A1 from Vip-IRES-Cre and Ndnf-IRES-Cre mice, respectively, whereas L4 Pyr neurons were recorded in both genotypes. **(B)** Example EPSCs evoked by minimal stimulation (V_hold_ = -70 mV) in L4 Pyr, L1 VIP-INs, and L1 NDNF-INs. Traces correspond to five successful trials (black) and five failures (gray). **(C)** Quantification of amplitude **(left)**, onset latency (**middle**), and rise time (**right**) of minimal stimulation-evoked EPSCs. **(D)** Example responses to trains of electrical stimulation applied to the thalamic axons at 20 Hz (V_hold_ = - 70 mV; mean ± SD of 5 trials). The recording procedure was the same as described in **(A)**, except that the electrical stimulus intensity was twice the threshold. **(E**,**F)** Amplitudes normalized to the EPSC amplitude evoked by the first electrical stimulus of the train (**left**) and ratio of the EPSC amplitudes in response to the last and first stimuli of the train (P_10_/P_1_, **right**) for each cell type during 10 Hz (**E**) and 20 Hz (**F**) repetitive stimulation. **(G)** Schematic of the experimental approach to record thalamic-evoked EPSCs with suprathreshold electrical stimuli to recruit intracortical inputs. Red dotted line represents the threshold to evoke EPSCs in response to electrical stimulation of thalamic axons. The recording procedure was as described in **(A)**, except that stimulation intensity was 5X the threshold. **(H)** Example evoked EPSCs in response to suprathreshold stimulation of the thalamic axons (V_hold_ = -70 mV; mean ± SD of 10 trials). **(I)** Quantification of peak amplitude (**left**), decay time constant (Τ_decay_, **middle**) and rise time (**right**). Stimulation artifacts were replaced by arrows for clarity in **B**, **D** and **H**. See **Supplementary Table 2** for details on statistical tests. *p < 0.05, **p < 0.01, ***p <0.001, ****p <0.0001.

### L1-INs show short-term depression of thalamic-evoked responses

Previous studies have shown that L4 Pyr neurons in ACx exhibit short-term depression in response to thalamic stimulation^87,90^, associated with synapses with a high initial probability of release^91^. This is in contrast to some IN subtypes in cortex that show facilitating thalamic inputs, suggesting low initial release probability of glutamate onto these neurons that is enhanced by repetitive stimulation^92,93^. Here, we analyzed the short-term plasticity of the thalamic responses in L1-INs by applying trains of electrical stimuli to the thalamocortical axons (**Figure 3D-F**). We found short-term depression for all cell types, as revealed by the significantly smaller EPSC amplitude in response to the last pulse in the train as compared to the first EPSC, both at 10 Hz and 20 Hz (**Figure 3E**,**F**, **left**, and **Supplementary Table 2**). We found no differences in the short-term depression across cell types, quantified as the ratio between the evoked EPSC amplitudes in response to the last and first stimulation pulses (**Figure 3E**,**F**, **right**, and **Supplementary Table 2**). Therefore, both L1 VIP- and NDNF-INs in A1 show robust short-term depression of thalamic-evoked responses, comparable to the characteristic depression observed in L4 Pyr neurons.

### Suprathreshold thalamic responses in L1-INs are smaller and slower than those recorded in L4 Pyr neurons

Finally, we aimed to understand how intracortical connectivity shapes the thalamic-driven responses of A1 L1-INs by using suprathreshold electrical stimulation to recruit both monosynaptic and disynaptic inputs (**Figure 3G**). Consistent with the strong recurrent connectivity of excitatory neurons in ACx^94,95^, L4 Pyr cells showed large suprathreshold EPSCs when thalamic axons were stimulated with an electrical current five times greater than the threshold stimulation intensity (**Figure 3H**). Both L1 VIP-INs and NDNF-INs showed significantly smaller suprathreshold EPSC amplitudes and slower EPSC decay kinetics as compared to L4 Pyr neurons (**Figure 3H**,**I** and **Supplementary Table 2**). Rise kinetics were similar for all three cell types under these recording conditions (**Figure 3H**,**I** and **Supplementary Table 2**). Together, these results highlight potential differences in the intracortical connectivity between L4 Pyr neurons and L1-INs in A1, indicating that the L1-INs are less coupled to the local cortical circuit.

## Discussion

Sensory cortices integrate information about specific features of the sensory environment with contextual signals that convey information about behavioral state, aversive and rewarding outcomes, and expectations^2,96^. The sites and mechanisms for this integration within the complex cortical circuits are not fully understood. Across sensory cortices, recent studies have highlighted the L1-INs as hubs for inputs from diverse sensory and non-sensory brain regions that modulate the activity and plasticity of deeper-layer excitatory neurons in an experience- and state-dependent manner^21,24,39,44,47,48,64–66,97^. However, thalamocortical synapses onto these L1-INs have not been fully characterized. In this study, we investigated the connectivity between the auditory thalamus and the two largest groups of L1-INs, VIP- and NDNF-INs^63^, within A1. Our results show that the majority of presynaptic thalamic neurons targeting these interneurons are located within the primary auditory thalamus, the MGBv. These L1-INs receive strong, monosynaptic connections from the MGBv with similar strength to the well-studied L4 Pyr neurons. Together, these results suggest that L1-INs receive robust inputs from neurons arising from the tonotopically-organized MGBv that convey information about the precise spectral and temporal features of sounds^5,56,60^.

We identify a robust pathway from the primary auditory thalamus (MGBv) directly to two major subtypes of L1-INs. These results are unexpected given that the canonical cortical model dictates that the primary sensory thalamic nuclei target the middle cortical layers (L3b and L4). Our retrograde tracing experiments show that the ipsilateral MGB forms monosynaptic connections with VIP- and NDNF-IN subpopulations in superficial ACx (**Figure 1**), consistent with previous reports showing thalamic inputs to inhibitory INs within the superficial layers of sensory cortices^39,44,46,72,98^. The MGB is composed of three major subdivisions: 1) the primary ventral division (MGBv) that contains neurons exhibiting short latency responses to sound, sharp frequency tuning and synchronized responses to amplitude modulated sounds, 2) the dorsal division (MGBd) that is composed of neurons that exhibit long-latency responses, broad, multi-peaked frequency tuning and less synchronization to amplitude modulations, and 3) the medial division that contains neurons with heterogeneous frequency tuning and modulation by non-auditory inputs^6,56,99^. Anatomical studies across species show that neurons arising from the MGBd generally target the superficial cortical layers^59,100,101^ and that large diameter axons from MGBm target L1^100^. Therefore, we were not surprised to find that both VIP- and NDNF-INs within superficial ACx receive inputs from the MGBd and MGBm. We also discovered a small input to these superficial INs from the multisensory PIN/PP regions in the auditory thalamus. This result is in agreement with recent studies showing higher-order thalamic inputs to NDNF-INs in auditory association areas^47^ and visual cortex^44^. The higher-order auditory thalamic regions receive multimodal inputs from diverse cortical and thalamic areas that receive somatosensory and visual information, non-sensory midbrain structures and neuromodulatory centers^47,52,55^. Projections from higher-order nuclei to L1 are prominent across sensory cortices, targeting both L1-INs and the apical dendrites of deeper layer excitatory neurons^47–51,102–105^, and are functionally relevant in learning and plasticity^47–49,53,104^.

Strikingly, the vast majority of presynaptic MGB neurons targeting the superficial VIP- and NDNF-INs were located in the primary MGBv. This primary thalamic nucleus receives the majority of its inputs from the inferior colliculus in the midbrain and sends tonotopically-organized projections to ACx^5,24,51,56,60^. These tufted neurons all belong to one morphological class of thalamic neurons that differ from the stellate neurons that are only located within the higher-order MGB subdivisions^99^. It has been shown across species that these MGBv neurons predominantly target L4 of A1^59,86,106^. More recently, however, it has been reported that MGBv neurons can also bifurcate in L4 and send collaterals towards superficial cortical layers^51^, also shown for first-order visual and somatosensory thalamic neurons^58,107,108^. This raises the possibility that the same frequency-tuned MGBv neurons may target both L4 Pyr and L1-INs within the same cortical column. However, it should be noted that in visual cortex, primary thalamic inputs may convey distinct sensory information to L1 and L4^34^. Our results motivate future *in vivo* studies that examine the auditory information that is conveyed by MGBV inputs to the L1-INs.

Our functional experiments revealed robust, monosynaptic thalamocortical connections onto L1 VIP- and NDNF-INs in A1 that were comparable in strength to those recorded in the L4 Pyr excitatory neurons (**Figure 2** and **Figure 3**). Moreover, the short-term depression of thalamic-evoked responses was similar between the L1-INs and L4 Pyr neurons, suggesting similar presynaptic release properties onto these cell types. This short-term depression of MGB-evoked responses also resembles that observed in parvalbumin (PV)-expressing INs^87,92,109^, which receive strong thalamic inputs across sensory cortex^57,92^ but differ from the somatostatin (SST)-expressing INs, which show short-term facilitation of thalamic-evoked inputs^92,93,109^. These results are consistent with anatomical studies showing that MGB neurons robustly innervate L1 within ACx^5,24,39,47,51,57,59^. Interestingly, only a subset of deeper-layer VIP-INs in ACx and primary somatosensory barrel cortex respond to stimulation of the MGBv^57^ or primary ventrobasal (VB) nucleus of the somatosensory thalamus^105^, respectively. Similarly, ∼65% of 5-HT_3A_R-expressing neurons, which include both VIP- and NDNF-IN subpopulations^63,110^, within L3/4 of the primary somatosensory cortex receive weak monosynaptic inputs from the VB^70^. VIP- and NDNF-INs are heterogeneous populations and these IN subtypes in the superficial cortex show distinct morphological properties as compared to those in deeper layers^63,75,111,112^. Therefore, VIP- and NDNF-INs in the outermost layer in the ACx may differ from those residing in the deeper layers and be uniquely positioned to receive strong sensory inputs.

Although the L1-INs and L4 Pyr neurons showed similar monosynaptic inputs from the MGB, large differences emerged between these cell types as the thalamic stimulation was increased to recruit recurrent intracortical activity. The currents recorded in L1 VIP- and NDNF-INs in response to suprathreshold thalamic stimulation were significantly more prolonged than those recorded in L4 Pyr excitatory neurons. It has been reported that MGBv inputs onto L4 Pyr are located along the perisomatic domain, whereas intracortical inputs are widely distributed across the dendritic arbor^106^. Although information about the subcellular distribution of thalamic and intracortical inputs onto VIP- and NDNF-INs is lacking, it is possible that the convoluted dendritic arbors that characterize these L1-INs^63,75,112^ affect the kinetics of responses originating from intracortical synapses on distal dendrites. Alternatively, the observed differences may arise from a different composition of glutamatergic receptor subtypes, which can differ between excitatory and inhibitory neuronal classes^113,114^. Future work that maps the thalamocortical and intracortical inputs onto subcellular domains of L1-INs may reveal the mechanisms underlying these observed differences in thalamic-driven responses.

Suprathreshold thalamic stimulation also evoked significantly smaller responses in L1 VIP- and NDNF-INs as compared to L4 Pyr excitatory neurons. Pyr neurons in ACx receive inputs across their dendritic arbors from local excitatory neurons^106^ that may dominate their spiking activity^115^ and shape their tuning to sensory stimuli^116–119^. Indeed, in our experiments suprathreshold thalamic stimuli evoked large EPSCs in Pyr cells, likely reflecting the excitatory inputs from L4, L2/3 and L6 Pyr neurons^94,95^. In contrast, L1 VIP- and NDNF-INs showed smaller EPSCs, suggesting that L1-INs are more disconnected from the local excitatory network^34^, although they receive inhibitory inputs from nearby interneurons^39,63,111^. These results highlight the function of L1-INs as integrators of unprocessed bottom-up sensory inputs with long-range top-down contextual inputs.

Due to their ability to integrate a wide range of inputs, VIP- and NDNF-INs in L1 have been described as sensors of global behavioral states and events, such as locomotion, attention, or reinforcement^34,62^. For this reason, most previous studies focused on inputs originating from subcortical regions like the cholinergic basal forebrain or the zona incerta^8,21,71,97^, multisensory thalamic regions^47,48,53,54,104^ and diverse sensory cortices^40,58,120^. However, the results presented here highlight that L1 VIP- and NDNF-INs in ACx are also vastly innervated by the primary auditory thalamus that conveys specific information about sound. Our results are consistent with previous results showing that L1-INs are activated by sensory stimuli^39^, and recent results from our laboratory showing that L1 VIP- and NDNF-INs exhibit tuning to simple and complex sound stimuli (personal communication, Sweeney, Thomas, Takesian). Together, these findings contribute to a growing shift in our view of these L1-INs towards ‘local circuit modulators’ rather than global broadcasters of behavioral states across cortex.

Our model of L1-INs as ‘local circuit modulators’ is consistent with recent findings that L1-INs send spatially precise axonal projections to specific cortical neuronal subtypes^24,39,44^. Although L1-INs produce the inhibitory neurotransmitter GABA, subsets of these interneurons have a specialized dis-inhibitory function in cortex— because they target other inhibitory interneurons, their activity leads to a net withdrawal of inhibition from glutamatergic neurons. NDNF-INs target PV interneurons, which reduces perisomatic inhibition onto deeper-layer Pyr neurons^44,74^ and VIP-INs target SST-INs, which reduces dendritic inhibition onto these Pyr neurons^66,67^. Both NDNF- and VIP-INs also directly inhibit the distal dendrites of Pyr neurons^39,47,74,121^. Disinhibiting L1-INs may directly project to the deeper layers to activate the cortical column, whereas other L1-INs project laterally to silence neighboring columns. In fact, *in vitro* and *in vivo* studies have demonstrated that these superficial interneurons increase the gain of cortical circuits^44,65,66^, punching ‘holes’ in the inhibitory networks below^67^. In the ACx, our results suggest that small groups of L1-INs within the same cortical column may be co-activated by frequency-tuned MGBv axons within the L1 tonotopic map^24,51^. Activation of these L1-IN ensembles may tune the sound-driven responses of the underlying thalamorecipient L4 Pyr neurons. New approaches for gain- and loss-of-function of small L1-IN ensembles may provide insight into their function in controlling the tuning and plasticity of deeper-layer excitatory neurons.

In addition to controlling moment-to-moment cortical activity, L1-INs are emerging as critical contributors to developmental and adult cortical plasticity and learning^24,39,48,64^. Indeed, both VIP- and NDNF-INs in ACx are activated by reinforcing stimuli, such as air puffs and footshocks, and are critical for auditory associative learning^39,66^. Interestingly, L1-INs may be sites of plasticity themselves^39,122,123^. For example, it has been shown that NDNF-INs undergo long-term changes in auditory-evoked responses following fear conditioning^39^. Thus, the MGBv and neuromodulatory synapses onto L1-INs may undergo heterosynaptic plasticity, causing long-term changes in how the L1-INs are recruited by sensory and non-sensory stimuli to dynamically modulate sensory-driven cortical activity. Our results open new avenues for future studies that will determine how distinct thalamic inputs to the L1-INs control their activity and plasticity during sound processing and learning.

## Acknowledgments

This work was supported by funds from the NIH NIDCD R01DC018353, and NIDCD F32DC018211 (to C.G.S.), the Nancy Lurie Marks Family Foundation, the Bertarelli Foundation, the Pew Latin American Fellows Program (to L.G.V.), the Herchel Smith Graduate Fellowship (to C.J.L.), the Science and Innovation Fellowship from Center on the Developing Child at Harvard University (to C.J.L.), the Amelia Peabody Scholarship (to C.J.L.). We thank Dr. Bernardo Sabatini (Harvard Medical School) and his lab members for kindly providing the SAD-ΔG-H2B-EGFP (RV-nEGFP) virus, the electrophysiology acquisition software, and for allowing us to use their slide scanner. We thank Dr. Daniel Polley (Mass Eye and Ear, Harvard Medical School) and his lab members for technical assistance and discussions, Dr. Maryse Thomas and Steven Minderler for feedback on the figures and manuscript, and Kasey Smith and Benjamin Glickman for animal care.

## Author Contributions

A.E.T. conceptualized this study. L.G.V. and A.E.T. designed the study. L.G.V. and C.J.L. performed the electrophysiology experiments and analyzed the electrophysiology data. C.P.M., L.G.V. and C.G.S. performed the surgeries for the rabies virus-based monosynaptic tracing, and C.P.M. processed the tissue for these experiments. C.P.M. and C.J.L. imaged and analyzed the rabies virus-based monosynaptic tracing data. L.G.V. and A.E.T. wrote the manuscript with input from all authors.

---

**Supplementary Figure 1.**
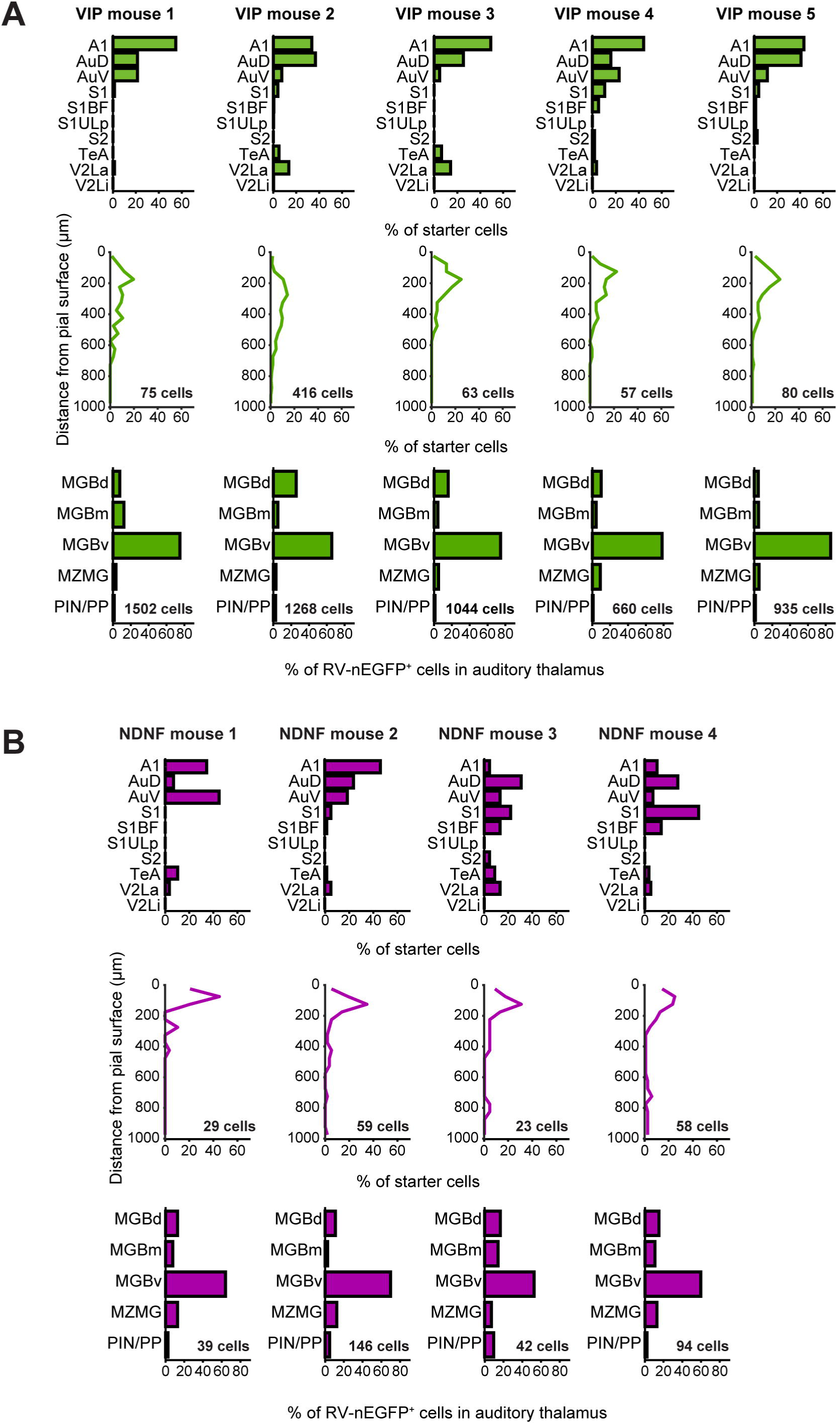
Individual experimental mouse data for rabies virus-based monosynaptic retrograde tracing. **(A**,**B)** Monosynaptic retrograde tracing data for five Vip-IRES-Cre mice (**A**, green) and four Ndnf-IRES-Cre (**B**, magenta). **Top.** Distribution of starter cells across cortical regions. **Middle**. Distribution of starter cells across cortical depth. Numbers indicate the total number of starter cells for each mouse. **Bottom**. Distribution of presynaptic RV-nEGFP^+^ cells across auditory thalamic regions. Numbers indicate the total number of presynaptic cells within auditory thalamic regions for each mouse. A1: primary auditory cortex; AuD: auditory dorsal area; AuV: auditory ventral area; S1: primary somatosensory cortex; S1BF: primary somatosensory cortex, barrel field; S1ULp: primary somatosensory cortex, upper lip region; S2: secondary somatosensory cortex; TeA: temporal association cortex; V2La: secondary visual cortex, lateral area, anterior region; V2Li: secondary visual cortex, lateral area, intermediate region; MGBv: ventral division of the medial geniculate body; MGBm: medial division of the medial geniculate body; MGBd: dorsal division of the medial geniculate body; MZMG: marginal zone of the medial geniculate body; PIN: posterior intralaminar nucleus; PP: peripeduncular nucleus.

**Supplementary Figure 2.**
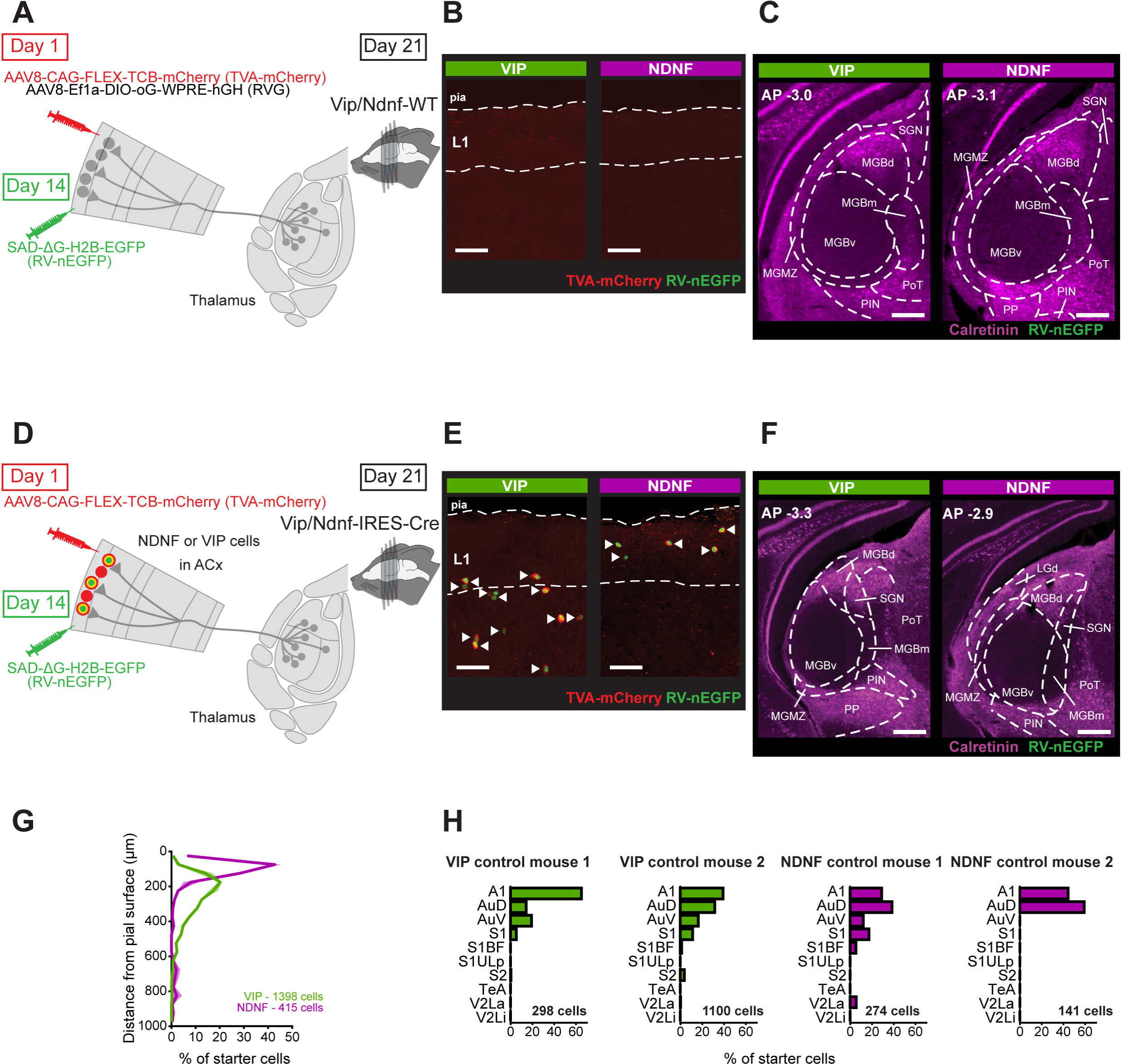
Rabies virus-based monosynaptic retrograde tracing validation. **(A)** Schematic of the approach to test the specificity of our starter cell populations for the monosynaptic retrograde tracing experiments. Vip/Ndnf-WT mice were injected with the same Cre-dependent viruses (TVA-mCherry and RVG) and RV-nEGFP used in the experimental mice, following the same procedure. This control was carried out in two Vip- and two Ndnf-WT mice. **(B)** Example confocal images of ACx regions of Vip-WT and Ndnf-WT mice, injected with Cre-dependent TVA-mCherry, Cre-dependent RVG, and RV-nEGFP. No starter cells were found in the ACx of Vip/Ndnf-WT, confirming the specificity of our Cre-dependent virus expressing TVA-mCherry. Scale bar: 50 µm. **(C)** Example epifluorescence images of auditory thalamic sections showing no RV-nEGFP^+^ neurons, consistent with the absence of starter cells in ACx. Calretinin immunostaining delineates the higher-order thalamic areas. **(D)** Schematic of the experimental approach to test for viral leak expression of RV-nEGFP. Vip- and Ndnf-IRES-Cre mice were injected with the Cre-dependent AAV expressing TVA-mCherry, omitting the injection of the Cre-dependent RVG required for monosynaptic retrograde infection. Subsequently, RV-nEGFP was injected following the same protocol employed in the experimental mice. **(E)** Example confocal images of ACx regions of Vip-IRES-Cre and Ndnf-IRES-Cre mice, injected with TVA-mCherry and RV-nEGFP. White arrows indicate representative starter cells in ACx, identified by the co-localization of mCherry and EGFP. Scale bar: 50 µm. **(F)** Example epifluorescence images of brain sections containing the auditory thalamic regions showing no RV-nEGFP^+^ neurons despite the presence of starter cells in ACx. Calretinin immunostaining delineates the higher-order auditory thalamus. **(G)** Distribution of starter cells across cortical depth in Vip/Ndnf-IRES-Cre control mice. Inset indicates the total number of starter cells in the cortex. Data corresponds to two mice of each genotype. **(H)** Distribution of starter cells in individual Vip-IRES-Cre (green) and Ndnf-IRES-Cre (magenta) mice used as controls to test for viral leak expression of RV-nEGFP. Inset indicates the total number of starter cells in each mouse.

## Methods

### Mice

Vip-IRES-Cre, Vip-WT, Ndnf-IRES-Cre, Ndnf-IRES-FlpO and Ai9 mice were purchased from Jackson Laboratories. Mice were bred in house and housed under standard laboratory conditions (12:12 hr light:dark cycle, access to food and water *ad libitum*). Both male and female adult (P60-90) mice were used for experiments. All experiments were approved by the Institutional Animal Care and Use Committee at Massachusetts Eye and Ear.

### Surgeries

Mice were deeply anesthetized with isoflurane 5% in O_2_ and moved to a stereotactic surgery rig, where isoflurane levels were lowered to 1.5%-2% and maintained throughout the procedure. At the start of the surgery, Buprenex (0.01 mg/ml; dosage 0.05 ml/10 g) and Meloxicam (0.2 mg/ml; dosage 0.1 ml/10 g) were administered via subcutaneous injections. After shaving and disinfecting the head, a small incision was made in the skin to expose the skull. The skull was leveled, and burr holes were made using a dental drill (MH170, Foredom Electric Co.) at the corresponding coordinates. For auditory cortex (ACx) surgeries, the head was rotated to the right to obtain a clear visual of the left ACx. All injections were performed using glass capillaries pulled with a micropipette puller (P-97, Sutter Instrument) and using a syringe (1701RN 10 µl, 80030, Hamilton Company) with a motorized piston (UltraMicroPump UMP3, World Precision Instruments) or a motorized programmable injector (Nanoject III, 3-000-207, Drummond Scientific Co.).

For electrophysiological recordings of monosynaptic primary thalamic inputs onto layer 1 (L1) VIP and NDNF interneurons (VIP-and NDNF-INs), and layer 4 Pyr (L4 Pyr) excitatory neurons, Vip-IRES-Cre x Nndf-IRES-FlpO mice were injected within the ventral portion of the left medial geniculate body (MGBv, in mm: AP: -3.12, ML: 2, DV: -3.1) with AAV2-CAG-ChR2-mCherry (1.55x10^12^; 100 nl at 20 nl/min). During the same surgical procedure, these mice were simultaneously injected with AAV8–CAG-FLEX-tdTomato and AAV1-CAG-fDIO-mNeonGreen (1.7x10^12^ and 9x10^12^, respectively; 500 nl at 20 nl/min) in two separate sites within the left ACx (in mm, AP = -2.0/-2.5, ML = 4.7, DV = -0.5, raising the glass capillary 0.1 mm every 100 nl injected).

For monosynaptic rabies virus tracing experiments, Ndnf-IRES-Cre or Vip-IRES-Cre mice were injected with Cre-dependent viruses expressing the avian EnvA receptor, TVA, fused to mCherry (TVA-mCherry) and the rabies virus glycoprotein (RVG) in the left ACx (in mm, AP = -2.3, ML = 4.7, DV = -0.1/-0.2; AAV8-CAG-FLEX-TCB-mCherry, 5.06x10^12^; AAV8-Ef1a-DIO-oG-WPRE-hGH, 1.24 x 10^13^; 200 nl in total at 20 nl/min). Two weeks later, a constitutive RVG-deleted, EnvA-pseudotyped rabies virus expressing nuclear-targeted EGFP (RV-nEGFP) was injected into ACx (Sabatini lab gift). Coordinates were the same as for Cre-dependent viruses and two injections were made at anteroposterior ends of the same burr hole (3.5x10^8^; 200 nl per site, at 20 nl/min). We validated our monosynaptic tracing strategy with control experiments. The first control confirmed that the starter neurons were restricted to Cre-expressing neurons by injecting all three viruses in WT mice (WT littermates, Ndnf-WT, or Vip-WT). We did not find any starter cells in ACx or presynaptic cells in auditory thalamic regions (**Supplementary Figure 2**A-C), confirming the requirement of the Cre recombinase to obtain starter cells. The second control evaluated potential leak expression of RV-nEGFP by omitting the injection of the virus that allows for RVG expression in Vip/Ndnf-IRES-Cre mice. Despite robust expression of starter cells in ACx, no presynaptic RV-nEGFP^+^ cells were observed in the thalamus (**Supplementary Figure 2**D-H).

### *In vitro* whole-cell recordings

Mice were anesthetized with isoflurane 5% in O_2_ and Fatal Plus administered intraperitoneally (18 mg/ml; dosage 0.1 ml/10 g). Immediately following anesthesia, mice were perfused with ice cold slicing artificial cerebrospinal fluid (ACSF) containing (in mM)^124^: 160 sucrose, 28 NaHCO_3_, 2.5 KCl, 1.25 NaH_2_PO_4_, 7.25 glucose, 20 HEPES, 3 Na-pyruvate, 3 Na-ascorbate, 7.5 MgCl_2_, 1 CaCl_2_. Following perfusion, mice were decapitated, the brain was quickly removed and thalamocortical slices (300 µm) obtained at an angle of 15° from horizontal view^86^ were collected in ice cold slicing ACSF on a vibrating microtome (Leica Microsystems; VT1200S). Slices were placed in a chamber for 30-35 minutes at 35° C in recovery ACSF, containing (in mM)^124^: 92 NaCl, 28.5 NaHCO_3_, 2.5 KCl, 1.2 NaH_2_PO_4_, 25 glucose, 20 HEPES, 3 Na-pyruvate, 5 Na-ascorbate, 4 MgCl_2_, 2 CaCl_2_. After recovery, slices were transferred to a chamber with recording ACSF, containing (in mM): 125 NaCl, 2.5 KCl, 1.25 NaH_2_PO_4_, 25 NaHCO_3_, 25 glucose, 1 MgCl_2_, 2 CaCl_2_^24^ and maintained for at least one hour at room temperature. Slices were moved to the recording chamber and maintained under continuous superfusion with recording ACSF at near-physiological temperature (31-33° C) throughout all recordings. All solutions were continuously bubbled with CO_2_-O_2_ (95%-5%).

Patch pipettes (3–5 MΩ) were obtained from borosilicate glass capillaries pulled with a micropipette puller (P-97, Sutter Instrument) and filled with voltage-clamp internal solution, containing (in mM)^125^: 100 KCl, 40 K-gluconate, 8 NaCl, 10 HEPES, 2 MgCl_2_, 0.1 EGTA, 2 Mg-ATP, 0.3 Na-GTP and 5 QX-314 (lidocaine); pH 7.2 with KOH. Data were acquired from cells with initial series resistance below 30 MΩ (60% compensation). Data were collected at a sampling rate of 10 kHz with a Multiclamp 700B amplifier (Molecular Devices), low-pass filtered at 3 kHz and digitized with a digital-to-analog converter board (National Instruments, NI-USB-6343). Custom-designed Matlab software was used for data acquisition and analysis (modified from Sabatini lab). All experiments were carried out in a motorized upright microscope (Scientifica, SliceScope Pro 1000) coupled to a CCD camera (Hamamatsu Photonics, Orca Flash 4.0 or Q Imaging, Retiga Electro). A subset of recorded cells was filled with biocytin 0.1-0.2% and slices were incubated with 4% PFA in 0.1 M phosphate buffer (PB) overnight. On the following day, slices were washed in phosphate buffer saline (PBS) and incubated in Alexa Fluor-488/700 Streptavidin for 2 hours to analyze the morphology of the filled neurons.

Optogenetic experiments to record MGBv monosynaptic inputs onto L1 VIP-INs, L1 NDNF-INs and L4 Pyr neurons were carried out in Vip-IRES-Cre x Ndnf-IRES-FlpO mice expressing ChR2 in the MGBv. To identify VIP- and NDNF-INs, we used Cre- and Flp-dependent reporters in ACx, while L4 Pyr neurons were identified by their characteristic morphology under IR-DIC. Light-evoked monosynaptic responses to a wide-field 470 nm LED pulse (CoolLED, pE-100; 5 ms, ∼14 mW/mm^2^) were recorded in voltage-clamp configuration (V_hold_ = -70 mV) in the presence of bath-applied 1 µM TTX and 1 mM 4-AP. Recordings were obtained from at least one cell from each subtype in every slice to allow for the normalization of evoked currents in L1 VIP- and NDNF-INs to those in L4 Pyr neurons. This controlled for variability in ChR2 expression across slices. Glutamatergic currents were abolished by 20 μM DNQX and 50 μM AP5.

For electrically-evoked synaptic responses, experiments were performed in Vip-IRES-Cre or Ndnf-IRES-Cre crossed with the Ai9 mouse line (Cre-dependent TdTomato), which allowed for the identification of L1 VIP-INs and NDNF-INs, respectively. L4 Pyr neurons were identified as described above. We recorded electrically-evoked excitatory postsynaptic currents (EPSCs) from L1 VIP-INs, L1 NDNF-INs and L4 Pyr cells in whole-cell voltage clamp configuration (V_hold_ = -70 mV). The thalamic axons were stimulated by placing a custom-made bipolar electrode (FHC, M21CEP(CT1)) on the white matter to deliver 0.5 ms current pulses using a constant current stimulus isolator (A.M.P.I., Iso-Flex). For each recorded cell, the electrical stimulation threshold was determined as the current intensity required to evoke EPSCs. Minimal stimulation experiments used a stimulation intensity near threshold to obtain a 50% failure rate^87^, and 50 trials were recorded for each cell at an interstimulus interval (ISI) of 10 seconds (s). During post-hoc analysis, a Kernel density estimation was performed on the distribution of minimal stimulation-evoked EPSC amplitudes, and the amplitude of the smallest peak was reported as the putative single axon response for a given neuron. For short-term plasticity experiments, electrical stimulation was slightly increased to twice the threshold determined for each cell, and 5 trials were acquired with an ISI of 30 seconds. To evaluate responses to suprathreshold thalamic stimulation, the electrical stimulus was increased to 5 times the threshold determined for each cell, and 10 trials were acquired with an ISI = 10 seconds. Optogenetically- and electrically-evoked EPSC recordings were obtained from VIP- and NDNF-INs residing exclusively in L1 below the L1/L2 boundary of the primary ACx (A1), estimated to be at around 100 µm^71^. For VIP- and NDNF-INs, the average depth of recorded neurons was 103.37 ± 28.79 µm, and 60.35 ± 22.52 µm, respectively.

### Immunohistochemistry

One week following the RV-nEGFP injections, mice were anesthetized with isoflurane 5% in O_2_ and Fatal Plus administered intraperitoneally (18 mg/ml; dosage 0.1 ml/10 g). Mice were subsequently perfused with 4% PFA in 0.1 M PB and brains were incubated in PBS with azide overnight. The following day, brains were washed (3x) in PBS and transferred to a 30% sucrose in PBS solution. Brains were incubated for 3 days or until they sunk to the bottom of the container. The brains were then removed from the solution, lightly patted dry to remove any droplets of solution, placed in a falcon tube and on dry ice until frozen. Full brains were sliced into coronal sections (50 µm) using a cryostat. Floating slices were maintained in PBS with azide at 4°C. Slices spanning the ACx region and the auditory thalamus were immunostained with anti-calretinin and anti-mCherry antibodies (both at 1:2000; secondaries were used at 1:500), treated with 60 nM DAPI and mounted. Images were acquired on a confocal microscope at 20X (Leica Microsystems, SP8) or on a slide scanner at 10X (Olympus, VS200).

### Image analysis

Quantification of monosynaptic rabies tracing experiments was performed using the NeuroInfo software. Serial images (10X) were converted to jp2 format and loaded into Assemble Serial Sections within the BrainMaker workflow. Individual slices were defined as sections through automatic detection of slice edges. Section order was manually reviewed and edited. Automated section alignment was performed using section shape and image features to yield accurate results. The alignment was manually reviewed and adjusted. Automatic cell detection was then performed using the Detect Cells pipeline, and manually reviewed. Cells were detected across all sections. RV-nEGFP-TVA-expressing cells were detected on the FITC and mCherry channels, respectively. The Colocalize Markers tool was used to identify neurons expressing both the fluorophore markers for RV-nEGFP- and TVA-mCherry-expressing cellswithin 7 μm distance). Sections were then aligned to an unified common coordinate framework atlas (YSK atlas)^78^ using the Section Registration workflow. Calibration was performed based on the same atlas. Manual registration adjustments were made until the sections aligned with the atlas overlay, using calretinin immunostaining to confirm the accuracy of our method. Data points were then exported by selecting Map Experimental Data to Atlas and selecting all planes to be mapped to all brain regions.

### Statistical analysis

Statistics were conducted using GraphPad Prism or Matlab. Electrophysiology data was analyzed by a Wilcoxon matched-pairs signed rank test, the Friedman non-parametric repeated measures ANOVA or the Kruskal-Wallis non-parametric ANOVA, followed by Dunn’s multiple comparisons test. Differences in cell counts across regions of the auditory thalamus were assessed using a Wilcoxon matched-pairs signed rank test. Data are represented as mean ± SEM and traces are represented as mean ± SD, unless otherwise noted. The number of cells (n) and mice used in this study are provided in **Supplementary Table 1** and **Supplementary Table 2**.

