## Supplementary Table 1 for "Primary auditory thalamus relays directly to cortical layer 1 interneurons"

|  |  | **VIP (5 mice)** | **NDNF (4 mice)** | **Statistical test and p value** |
| --- | --- | --- | --- | --- |
| % of presynaptic cells in auditory thalamic regions - **Figure 1E** | MGBd | 12.55 ± 3.65 | 13.83 ± 1.24 | Wilcoxon rank sum test: W = 23, p = 0.56 |
|  | MGBm | 6.08 ± 1.57 | 8.84 ± 2.44 | Wilcoxon rank sum test: W = 23, p = 0.56 |
|  | MGBv | 74.93 ± 3.20 | 61.32 ± 3.56 | Wilcoxon rank sum test: W = 11, p = 0.032 |
|  | MZMG | 5.05 ± 0.99 | 11.26 ± 1.38 | Wilcoxon rank sum test: W = 29, p = 0.032 |
|  | PIN/PP | 1.39 ± 0.31 | 4.75 ± 1.69 | Wilcoxon rank sum test: W = 29, p = 0.032 |
