## Supplementary Table 2 for "Primary auditory thalamus relays directly to cortical layer 1 interneurons"

|  | **L4 Pyr** | **VIP** | **NDNF** | **Statistical test and p value** |
| --- | --- | --- | --- | --- |
| Peak amplitude (TTX+ 4AP) - **Figure 2E** | 300.00 ± 65.41 pA (n = 12 cells, 6 mice) | 280.00 ± 39.77 pA (n = 14 cells, 6 mice) | 228.60 ± 55.15 pA (n = 12 cells, 6 mice) | Kruskal-Wallis test with Dunn’s multiple comparisons test: H[2] = 1.58, p = 0.45 |
| Onset latency  (TTX + 4AP) - **Figure 2E** | 1.93 ± 0.12 ms (n = 12 cells, 6 mice) | 1.79 ± 0.08 ms (n = 14 cells, 6 mice) | 1.85 ± 0.07 ms (n = 12 cells, 6 mice) | Kruskal-Wallis test with Dunn’s multiple comparisons test: H[2] = 0.80, p = 0.67. |
| Normalized amplitude (TTX + 4AP) - **Figure 2E** | 1.00 ± 0.22 (n = 12 slices, 6 mice) | 1.51 ± 0.42 (n = 12 slices, 6 mice) | 1.23 ± 0.42 (n = 12 slices, 6 mice) | Friedman’s test with Dunn’s multiple comparisons test: Q = 0.50, p = 0.78 |
| Peak amplitude - **Figure 3C** | 26.13 ± 1.92 pA (n = 12 cells, 4 mice) | 24.60 ± 1.66 pA (n = 17 cells, 6 mice) | 22.79 ± 0.90 pA (n = 23 cells, 9 mice) | Kruskal-Wallis test with Dunn’s multiple comparisons test: H[2] = 2.25, p = 0.32 |
| Onset latency - **Figure 3C** | 8.54 ± 0.77 ms (n = 12 cells, 4 mice) | 12.00 ± 0.73 ms (n = 17 cells, 6 mice) | 11.50 ± 0.54 ms (n = 23 cells, 9 mice) | Kruskal-Wallis test with Dunn’s multiple comparisons test: H[2] = 9.82, p = 0.0074 |
| Rise time - **Figure 3C** | 3.16 ± 0.29 ms (n = 12 cells, 4 mice) | 3.19 ± 0.29 ms (n = 17 cells, 6 mice) | 2.91 ± 0.21 ms (n = 23 cells, 9 mice) | Kruskal-Wallis test with Dunn’s multiple comparisons test: H[2] = 0.66, p = 0.72 |
| Normalized amplitude, 10 Hz - **Figure 3E** | P_1_ = 1.00 ± 0.29 - P_10_ = 0.58 ± 0.21 (n = 11 cells, 4 mice) | P_1_ = 1.00 ± 0.27 - P_10_ = 0.47 ± 0.11 (n = 8 cells, 4 mice) | P_1_ = 1.00 ± 0.28 - P_10_ = 0.60 ± 0.16 (n = 10 cells, 6 mice) | Friedman's test with Dunn’s multiple comparisons test - L4 Pyr: Χ^2^[9] = 75.50, p(1 vs 10) < 0.0001 - VIP: Χ^2^[9] = 34.53, p(1 vs 10) = 0.0002 - NDNF: Χ^2^[9] = 42.09, p(1 vs 10) = 0.0002 |
| P_10_/P_1_, 10 Hz - **Figure 3E** | 0.49 ± 0.05 (n = 11 cells, 4 mice) | 0.61 ± 0.12 (n = 8 cells, 4 mice) | 0.69 ± 0.10 (n = 10 cells, 6 mice) | Kruskal-Wallis test with Dunn’s multiple comparisons test: H[2] = 2.32, p = 0.31 |
| Normalized amplitude, 20 Hz - **Figure 3F** | P_1_ = 1.00 ± 0.28, P_10_ = 0.44 ± 0.15 (n = 9 cells, 4 mice) | P_1_ = 1.00 ± 0.25, P_10_ = 0.27 ± 0.04 (n = 9 cells, 4 mice) | P_1_ = 1.00 ± 0.31, P_10_ = 0.38 ± 0.10 (n = 11 cells, 6 mice) | Friedman's test with Dunn’s multiple comparisons test - L4 Pyr: Χ^2^[9] = 73.82, p(1 vs 10) < 0.0001 - VIP: Χ^2^[9] = 62.89, p(1 vs 10) < 0.0001 - NDNF: Χ^2^[9] = 51.28, p(1 vs 10) = 0.0003 |
| P_10_/P_1_, 20 Hz - **Figure 3F** | 0.39 ± 0.04 (n = 9 cells, 4 mice) | 0.40 ± 0.09 (n = 9 cells, 4 mice) | 0.60 ± 0.14 (n = 11 cells, 6 mice) | Kruskal-Wallis test with Dunn’s multiple comparisons test: H[2] = 2.44, p = 0.29 |
| Peak amplitude - **Figure 3I** | 652.60 ± 101.20 pA (n = 9 cells, 4 mice) | 232.00 ± 42.56 pA (n = 7 cells, 5 mice) | 356.00 ± 81.91 pA (n = 17 cells, 7 mice) | Kruskal-Wallis test with Dunn’s multiple comparisons test: H[2] = 8.352, p = 0.0154 |
| Τ_decay_ - **Figure 3I** | 27.81 ± 5.45 ms (n = 9 cells, 4 mice) | 68.28 ± 12.13 ms (n = 7 cells, 5 mice) | NDNF: 81.37 ± 10.28 ms (n = 17 cells, 7 mice) | Kruskal-Wallis test with Dunn’s multiple comparisons test: H[2] = 11.69, p = 0.0029 |
| Rise time - **Figure 3I** | 7.43 ± 0.81 ms (n = 9 cells, 4 mice) | VIP: 8.59 ± 1.43 ms (n = 7 cells, 5 mice) | 10.73 ± 1.54 ms (n = 17 cells, 7 mice) | Kruskal-Wallis test with Dunn’s multiple comparisons test: H[2] = 1.03, p = 0.58 |
